# Feedback control promotes synchronisation of the cell-cycle across a population of yeast cells

**DOI:** 10.1101/590844

**Authors:** Giansimone Perrino, Davide Fiore, Sara Napolitano, Francesca Galdi, Antonella La Regina, Mario di Bernardo, Diego di Bernardo

## Abstract

The periodic process of cell replication by division, known as cell-cycle, is a natural phenomenon occurring asynchronously in any cell population. Here, we consider the problem of synchronising cell-cycles across a population of yeast cells grown in a microfluidics device. Cells were engineered to reset their cell-cycle in response to low methionine levels. Automated syringes enable changing methionine levels (control input) in the microfluidics device. However, the control input resets only those cells that are in a specific phase of the cell-cycle (G1 phase), while the others continue to cycle unperturbed. We devised a simplified dynamical model of the cell-cycle, inferred its parameters from experimental data and then designed two control strategies: (i) an open-loop controller based on the application of periodic stimuli; (ii) a closed-loop model predictive controller (MPC) that selects the sequence of control stimuli which maximises a synchronisation index. Both the proposed control strategies were validated in-silico, together with experimental validation of the open-loop strategy.

## I. Introduction

Recently, applications of Control Engineering to steer biological processes in living cells, also known as Cybergenetics, have been demonstrated [1]–[7]. These applications are based on innovative technological platforms exploiting microfluidics and optogenetics to implement real-time feedback control, where the controller is implemented as a software in the computer.

Here, we investigated the possibility to synchronise a population of budding yeast by forcing cells to divide in a coordinated fashion for multiple generations. During its lifespan, the eukaryotic cell undergoes a sequence of cyclically repeated steps that lead to its division via a process known as the cell-cycle. It can be divided into four main phases as shown in Figure 1: G1 (growth phase), S (DNA synthesis starts), G2 (growth phase) and M (mitosis) that includes the actual cell division. In budding yeast, the three endogenous genes encoding the G1 cyclins (*CLN1, CLN2* and *CLN3*) are expressed in the late G1 phase and promote the transition from G1 to the S phase. Moreover, expression of just one of these three cyclins is sufficient for the G1 to S transition to occur.

**Fig. 1.**
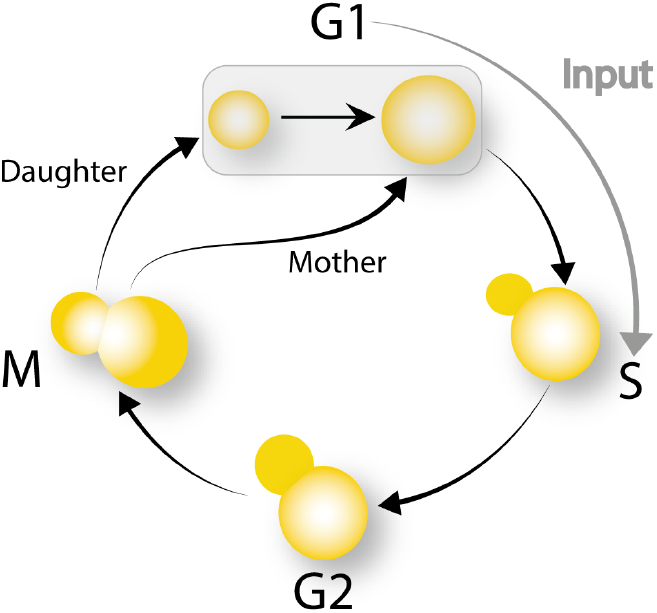
Cell-cycle process in a budding yeast manipulated genetically to reset the cell-cycle from the G1 phase to the S phase in the presence of an external stimulus [8].

Biologists have developed different methods to force a cell population to divide synchronously. However, all of these methods do not dynamically synchronise the cell population, but essentially they just force each cell in the population to start from the same initial condition. Therefore, after a few generations, the population de-synchronises because of environmental and cell specific disturbances.

To address the cell-cycle synchronisation problem from a Control Engineering point of view, we chose a budding yeast strain where the endogenous *CLN3* gene was deleted while the methionine-repressible promoter P_*MET3*_ was inserted upstream of the endogenous *CLN2* gene [8]. The resulting strain will thus not express *CLN2* if grown in a methionine-rich medium, but it will be able to cycle nevertheless, as the endogenous *CLN1* is still present in this strain. However, by switching growth medium from methionine-rich to methionine-depleted medium, the cell can be forced to transition from the G1 phase to the S phase thanks to the resulting expression of *CLN2*, as represented in Figure 1. Hence the growth medium can be used as a control input.

To track the cell-cycle progression in time, a yellow fluo-rescent protein (YFP) was fused to the endogenous *CDC10* gene, which encodes a protein expressed in the M phase during cell division at the edge between the mother and daughter cell (bud neck). The intensity of the YFP represents the system output.

We derived a simple phase-oscillator model of the cell-cycle in this strain, following a modelling framework described in Perrino *et al.* [9], whose parameters were set by considering experiments reported by Charvin *et al.* [8]. We used this model both for numerical simulations and for designing control strategies. In order to measure the synchronisation level, we considered a performance index based on Kuramoto order parameter. We then designed an open-loop control strategy using a periodic external input, as previously propose by Charvin *et al.* [8]. Next, we implemented a model predictive control (MPC) strategy to check whether a feedback control strategy could improve the performances as compared to the open-loop strategy.

Finally, we performed a preliminary experimental validation of our control strategies exploiting a microfluidics-based feedback control platform that we have recently developed [7], [10].

Our results show that the results presented in our previous work can be reached also considering cells able to cycle with a their own dynamics.

## II. Model derivation and identification

To model the cell-cycle, we exploited the concept of phase reduction [11], [12]. We assumed that the cell-cycle can be described as a dynamical system of the general form:

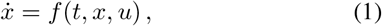

where *t* ∈ ℝ is time, *x* ∈ ℝ^*n*^ is the state vector and *u* ∈ ℝ is the control input. If (1) has an exponentially stable limit cycle *γ* ⊂ ℝ^*n*^ with period *T_d_*, then (1) is an oscillator that, according the phase reduction method, can be modelled as a dynamical phase oscillator

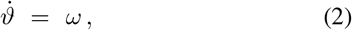

where *ϑ* is the phase of the oscillator on the unit circle 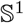 and 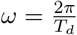 is the angular frequency.

Let *ϑ_c_* be the cell-cycle phase at budding, i.e. the phase at which the G1 to S transition occurs. As the cell-cycle is coupled to cell growth [8], the phase dynamics can be described as:

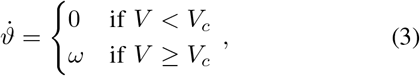

where *V* is the cell volume, and *V_c_* is the critical volume, defined as the minimal volume necessary for cell-cycle progression. The phase linearly increases with rate *ω*, unless the cell volume is less than the critical value *V_c_*. To model volume dynamics, we assumed that a cell grows exponentially only during the G1 phase, thus neglecting the mass generated during the S-G2-M phases, most of which is transferred to the growing bud (daughter cell) [8]:

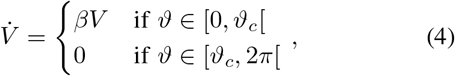

where *β* > 0 is the volume growth rate. Finally, we assume that at *ϑ* = 2*π* cell division occurs and the bud detaches from the mother (M) cell, and thus forming a new daughter (D) cell. Let *ϑ_M_* be the mother cell phase, *V_M_* its volume; *ϑ_D_* the daughter cell phase and *V_D_* its volume. At each division, the birth of a daughter cell is modelled according to the following rule:

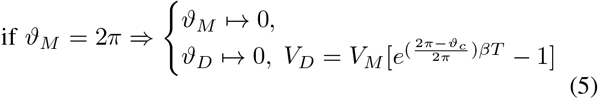

where at division, *V_D_* is equal to the volume growth occurred during the S-G2-M phases at bud level.

Let *u* ∈ {0,1} be the external trigger input to the system with *u* =1 corresponding to methionine-depleted medium and *u* = 0 to methionine-rich medium. The effect of the input can be modelled with the following reset rule:

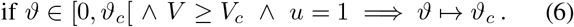

According to this rule, the control input *u* will force the G1 to S transition only in those cells that are in the G1 phase and have a volume *V* > *V_c_*, while the other cells will continue to cycle unperturbed. This constraint is a major limitation to the performance of any controller.

The numerical values of the parameters of this cell-cycle model were inferred from experimental data presented in Charvin *et al*. [8]. Thus, the cell-cycle period has been set to 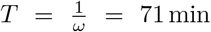, the volume growth rate to *β* = 0.0083 [a.u.]min^-1^, and the critical volume to *V_c_* = 1 [a.u.]. Moreover, the cell-cycle phase at budding has been set to 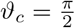.

## III. Problem statement

Let 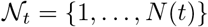 be the finite set of all cells in the population at time *t*, where 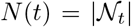 is the number of cells in the population at time t. Note that the number of cells may vary in time as a consequence of cell births and deaths. However, here we consider only cell birth events, therefore *N*(*t*): *t* → ℕ will be at most a nondecreasing function.

Consider each cell 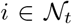 as an individual agent whose cell-cycle progression is mathematically described by the model composed by (3) and (4), together with the reset rule (6) and the division rule (5).

The control objective is to synchronise the cell-cycle across the cell population. Synchronisation in a population of oscillators can be quantified by means of the Kuramoto order parameter *R* [12] defined as the magnitude of the complex number:

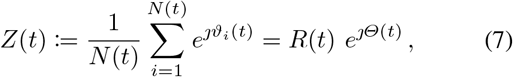

where *ϑ_i_* is the phase of cell 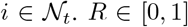 represents the mean phase coherence, an index to evaluate the synchronisation among a population of oscillators. When *R* is equal to 1, all cells are synchronised onto the same phase.

To evaluate the performance of the control algorithms in synchronising the cell population, we introduce the cost function:

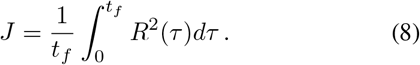

Thus, when *J* equals one then all cells are synchronised over the whole time interval.

As newborn cells may have an initial volume smaller than *V_c_*, it follows from (3) that their phases will remain equal to 0 for a certain time interval. Moreover, at this stage, these cells will not respond to the control input, hence taking into account their phases in (7) may result in misleading values of the control performance. To overcome this problem, we redefined *Z* (*t*) and *J*(*t*) by omitting non-cycling cells in their evaluation. Let 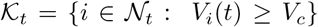 be the subset of cells whose volume is greater than the critical one at time *t*. Then, the Kuramoto order parameter associated to the cycling cells is defined as:

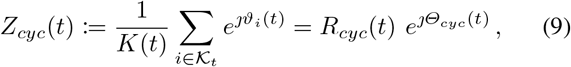

where *K*(*t*) is the cardinality of the set 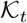 at time *t*. *R_cyc_* ∈ [0,1] is the mean phase coherence of the cycling cells. In the same way, we introduce the cost function associated to the synchronisation of the cycling cells, defined as:

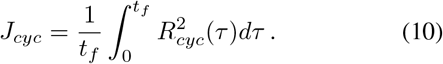

We will test control strategies in two different scenarios: (1) *constant population* where all cells are mother cells (i.e. *V_i_*(0) ≥ *V_c_*) and daughter cells are not considered so that the number of the cells is constant for all *t*. obviously, in this case, since every cell is a cycling mother we have that 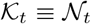 for all *t* and therefore *J_cyc_* = *J*; (2) *time-varying population* in which we also consider birth events of daughter cells. Thus two control problems can be formulated:

*Problem 1:* Given a *constant population* of N cells, whose dynamics is described by (3)-(4) with the phase reset law (6), compute the control input u that maximises *J_cyc_*.

*Problem 2:* Given a *time-varying population*, with an initial number of *N*(0) = *N*_0_ cells, whose dynamics is described by equations (3)–(5) with the phase reset law (6), compute the control input u that maximises *J_cyc_*.

### A. Numerical simulations

All in-silico simulations were performed using the MAT-LAB `odel5s` solver with event detection routines to accurately detect cell division at *ϑ_i_* = 2*π* and the critical volume *V_i_* = *V_c_* for cells with initial volume *V_i_* < *V_c_*.

For the constant population (Problem 1), we considered *N* = 100 mother cells, with an initial volume *V_i_*(0) for all the cells equals to the critical value *V_c_*. Instead, the initial phases *ϑ_i_* (0) of individual cells were uniformly spaced in the interval [0,2 *π*[.

For the time-varying population (Problem 2), we considered an initial number of *N*_0_ = 1 cell having initial volume *V*_1_(0) = *V_c_* and phase *ϑ*_1_(0) = *ϑ_c_*.

All simulations were run with the numerical parameters reported in Section II, for a time interval lasting *t_f_* = 15 *T*. Since we considered the same numerical parameters for each cell agent, the cell-to-cell variability across the cell population inherently arises from the cell division events.

## IV. Open-loop control

We designed a simple open-loop control strategy which consists in forcing the cells with a periodic control input of constant period *τ*. This strategy relies on the phase-locking phenomenon observed to occur in yeast when forced with a periodic external stimulus [8]. Numerical simulations for the control problems 1 and 2 (refer to Section III-A) were performed for different values of forcing period τ from 1 min to 142 min (that is two times the nominal period *T*) with increments of 1 min. As an example, we report in Figure 2 the response of the time-varying cell population, with initial number *N*_0_ = 1, to a control input of period *τ* = 64 min. In this case, the population exhibits an intermediate level of synchronisation as indicated by the evolution of the performance index *R_cyc_*(*t*). In Figure 5, we report the value of the cost indices *J* and *J_cyc_* as a function of the forcing period *τ* for both control problems.

**Fig. 2.**
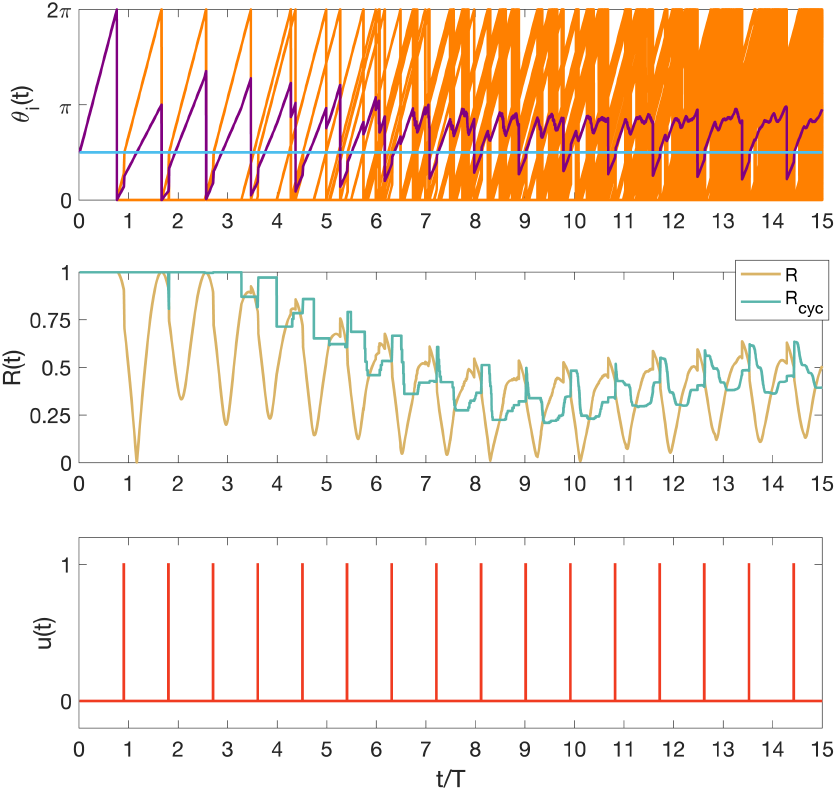
Example of an open-loop control simulation. Upper panel, time evolution of *ϑ_i_* (*t*), as well as their average value over the whole population, are reported respectively in orange and purple colours. Middle panel, time evolution of *R*(*t*) and *R_cyc_*(*t*) are reported respectively in yellow and green colours. Bottom panel, the corresponding control signal *u*(*t*), generated by the open-loop control assuming a forcing period *τ* = 64min, is reported in red colour.

It is apparent that the open-loop control strategy has a satisfactory performance for the constant population but clearly fails in the case of the time-varying population, as shown in Figure 3(b).

**Fig. 3.**
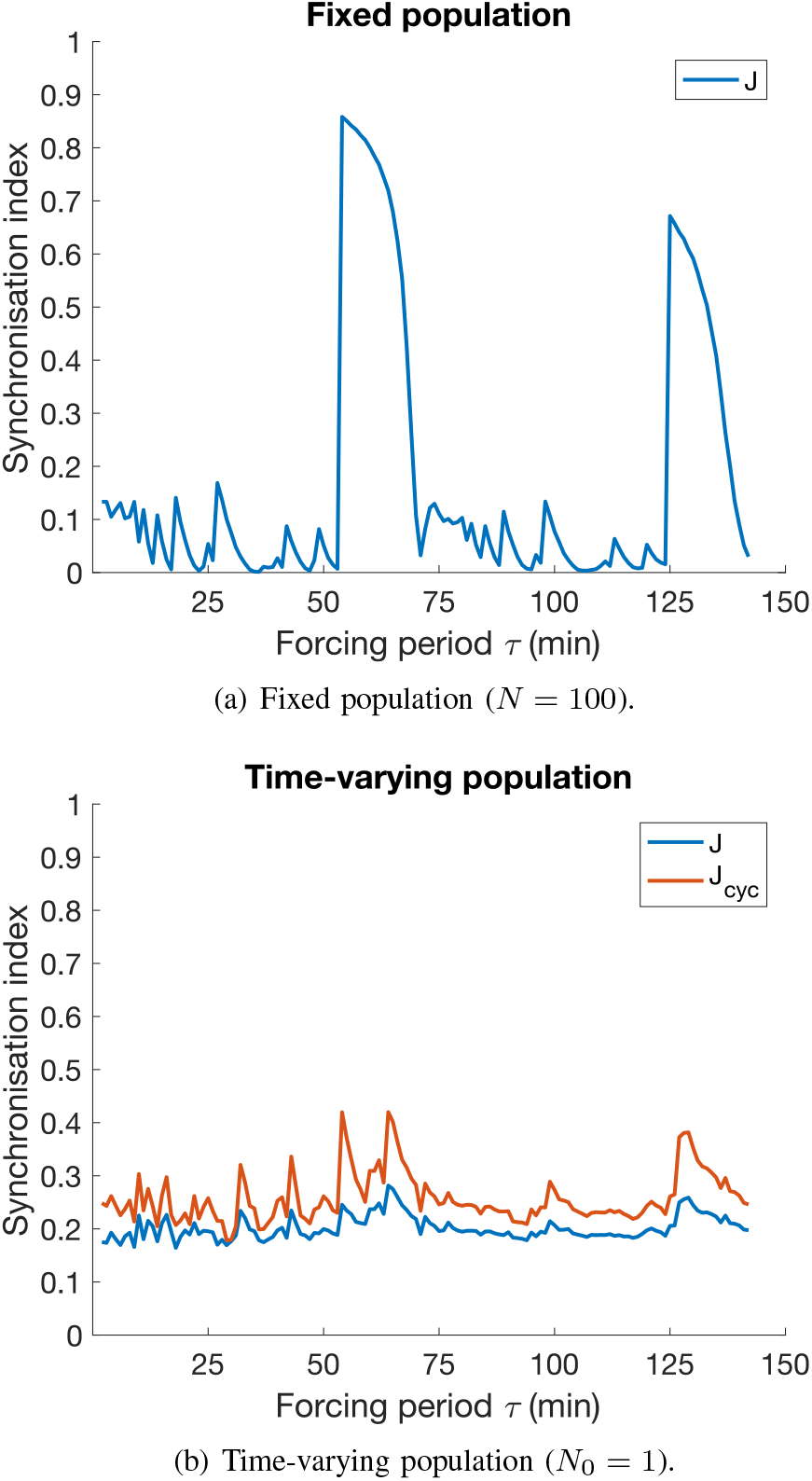
Open-loop control: synchronisation index as a function of the forcing period *τ. J* is the synchronisation index associated to the whole population, whereas *J_cyc_* is the synchronisation index computed considering only the cycling cells. *τ* varies from 1 min to 142 min (that is equal to 2T) with increments of 1 min.

## V. FEEDBACK CONTROL

Although being very simple, the previous open-loop control strategy requires precise tuning of the control parameter *τ* and has the drawback of not being robust to uncertainties and variation of the parameters of the cell-cycle model. To overcome these limitations, we devised a *model predictive control* algorithm, that is a feedback control strategy that can guarantee better performances in synchronising cell-cycle phases [9].

### A. Model Predictive Control

The MPC algorithm consists in solving an open-loop optimal control problem repeatedly over a receding horizon [13]. This means that at each iteration of the algorithm, the solution to the optimal open-loop control problem gives an optimal control input that minimises (or eventually maximises) a cost function over a finite prediction horizon *T_p_*. The optimal control input is applied only over a finite control horizon *T_c_* ≤ *T_p_*. Then, the optimisation is repeated again.

To address the problem of synchronising the cell-cycle phases, we chose as cost function the performance index *J_cyc_*, by setting *t_f_* = *T_p_* in equation (10). Since cell-cycle phases are synchronised when *J_cyc_* is equal to one, in this case we have to solve a maximisation optimal control problem.

To reduce the computational complexity of the optimal control problem, the control input 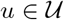 is assumed to be a finite sequence of triggers. Defining *P* £ ℕ as the maximum number of triggers that may be applied in a finite prediction horizon (*t, t* + *T_p_*], then the time interval occurring between two consecutive triggers is 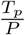. The feasible set 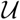 is defined as the set of all admissible control sequences. Considering only the finite prediction horizon *T_p_*, the set 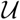 is composed by 2^*P*^ possible combinations of triggers. Thus, the optimal control problem is solved by maximising the performance index *J_cyc_* over the prediction horizon *T_p_*. The optimisation is achieved by exploring all the possible combinations of sequence of triggers.

For the numerical analysis, we set the length of both the prediction horizon *T_p_* and the control horizon *T_c_* equal to the nominal cell-cycle duration *T*, and the number of triggers *P* spanning in the range [1, 6]. Figure 4 shows the response of the time-varying cell population, with an initial cell number *N*_0_ = 1, to the control input generated by the MPC algorithm with control parameter *P* = 5.

**Fig. 4.**
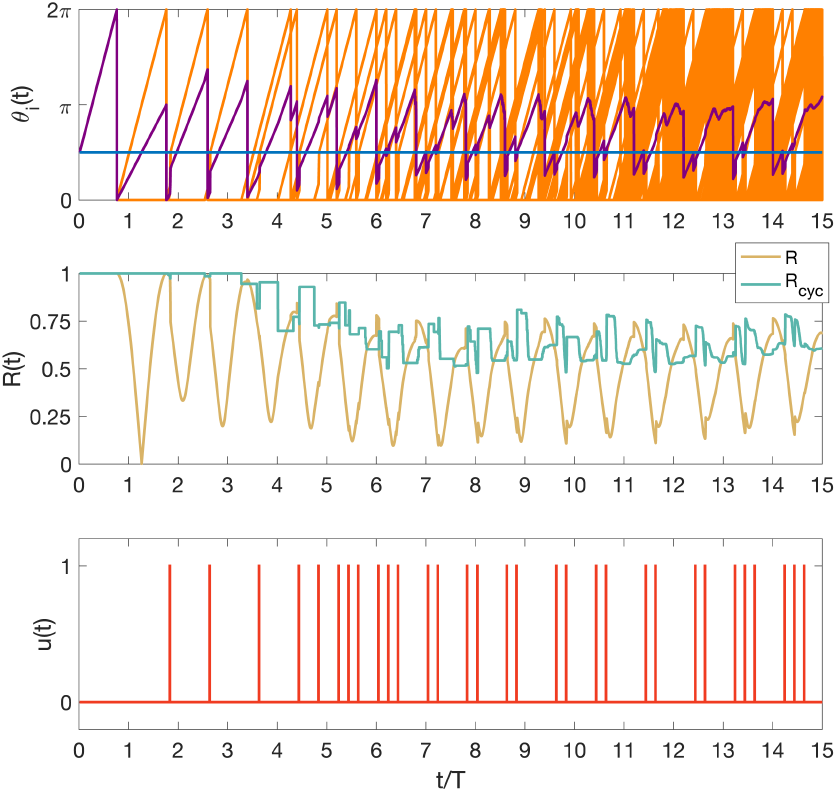
Example of a feedback control simulation using MPC. Upper panel, time evolution of *ϑ_i_*(*t*), as well as their average value over the whole population, are reported respectively in orange and purple colours. Middle panel, time evolution of *R*(*t*) and *R_cyc_*(*t*) are reported respectively in yellow and green colours. Bottom panel, the corresponding control signal *u*(*t*), generated by the MPC with control parameter *P* = 5, is reported in red colour.

In Figure 5 we report the value of the cost functions *J* and *J_cyc_* for the parameter *P* ranging from 1 to 6. The best performance are obtained for *P* equal to 5 and 6 in both fixed and time-varying population scenarios.

**Fig. 5.**
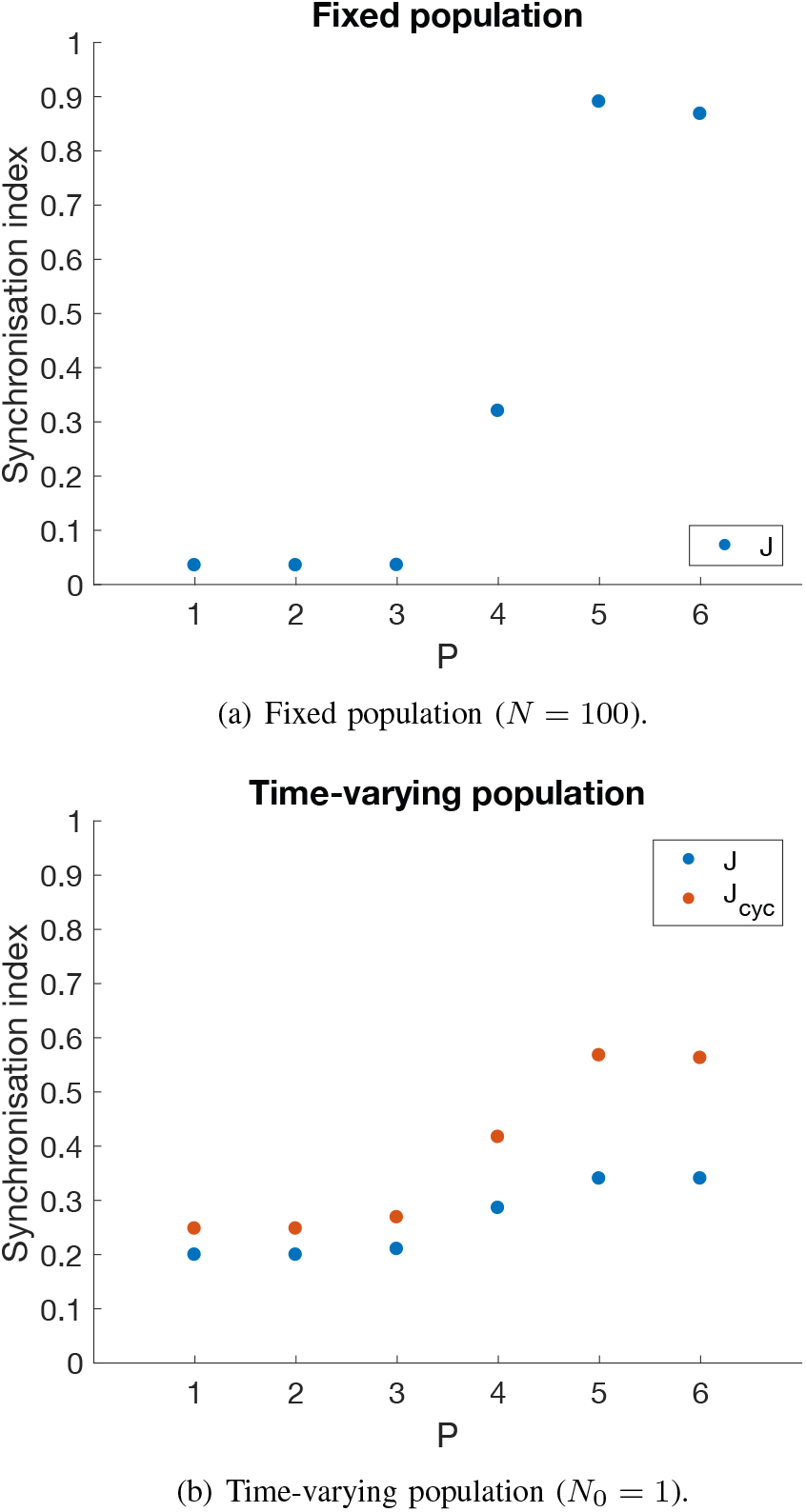
Model Predictive Control: synchronisation index as a function of the maximum number of triggers *P* that may be applied in a finite prediction horizon *T_p_*. *J* is the synchronisation index associated to the whole population, whereas *J_cyc_* is the synchronisation index computed considering only the cycling cells.

With respect to the synchronisation indices obtained with the open-loop control reported in Figure 3, the MPC has clearly better performances in both constant population scenario and especially in the time-varying population scenarios. Moreover, the MPC is obviously more robust to uncertainties in the model parameters since it does not require any tuning of parameters as in the open-loop controller. However, the performance of the MPC is strongly dependent on the particular choice of admissible control sequences U that have to be carefully selected.

## VI. Preliminary in vivo validation

To investigate the cell-cycle synchronisation problem experimentally, we performed a preliminary validation of the open-loop control strategy described in Section IV. To this end, we took advantage of a microfluidics-based experimental platform that was already used to control gene expression levels on living yeast cells [7], [10], [14]. We performed two experiments: (i) a calibration experiment where yeast cells were grown in methionine-rich growth medium (control input *u* = 0); (ii) an *open-loop control* experiment in which the growth medium is periodically switched between methionine-rich and methionine-poor with a period *τ* = 60 min (control input), i.e. close to the optimal 54 min period found with the numerical simulations (Figure 3). In this latter experiment, differently from the numerical simulations where the control input consisted of instantaneous pulses, the control input *u* is a square wave with a period *τ* = 60 min and a duty cycle *D* = 33%, that is methionine-poor medium was applied to the cell population for a duration of 20 min, that is one-third of the forcing period *τ* used in this experiment. This is necessary, as a shorter stimulus will not be sensed by the cell [8]. Both experiments lasted 14 hours, that are almost 12 times the nominal period of the cell-cycle of our yeast strain. The results of these two experiments are reported in Figure 6, where the output of each cell, i.e. the yellow fluorescence signal, is shown. Cells in the unforced experiment (Figure 6(a)) appear to be poorly synchronised when compared to those subjected to periodic stimuli (Figure 6(b)). Indeed, in the open-loop control experiment cells appear to be phase-locked with the external periodic stimulus (ratio 2:1). Nevertheless, not all of the cells appear to respond to every external stimuli. Further experiments, including closed-loop feedback control will need to be performed to validate the numerical simulations.

**Fig. 6.**
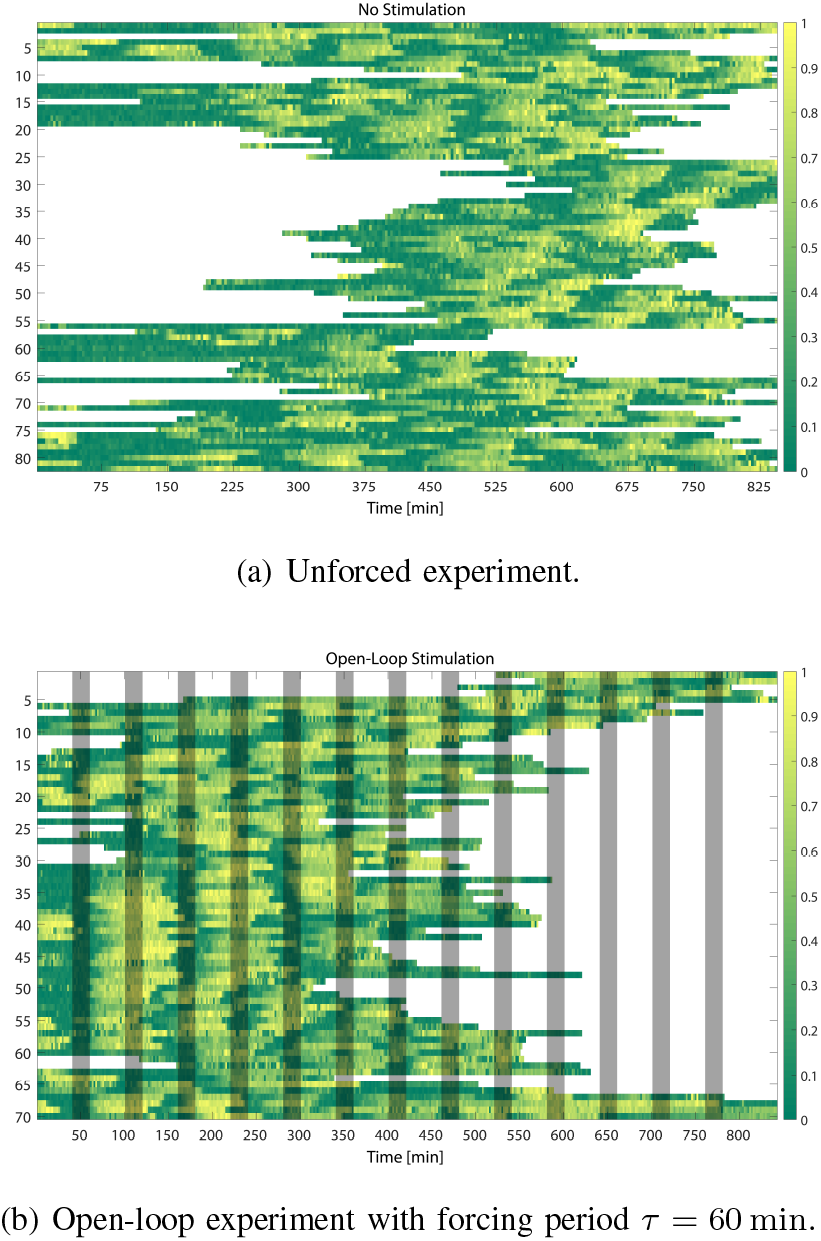
Open-loop experiments: single-cell traces of the yellow fluorescence reporter measured at single-cell level during two time-lapse open-loop experiments. Each single-cell trace was normalised between the maximum and minimum levels of fluorescence measured during the cell time-course. Then, all the single-cell traces were sorted with a single-linkage clustering algorithm. The duration of each signal is different as cells are being born and pushed out of the field of view during the course of the experiments. (a) Open-loop experiment in which no external stimulus was provided to the yeast cells. (b) Open-loop control experiment in which a periodic stimulus was provided to the yeast cells. In the latter case, the forcing period *τ* was equal to 60 min. Bud formation is highlighted in the heat maps by high values of the fluorescence signal, that is when the colours are close to yellow. Grey bars denote the external stimuli.

## VII. Conclusions

In this work, we have addressed the cell-cycle synchronisation problem in budding yeast considering a real biological scenario. We considered a budding yeast strain whose cell-cycle can be reset from the G1 phase to the S phase in the presence of a proper external stimulus, that is the absence of methionine in the growth medium supplied to the cells. We characterised the biological system deriving a phase-oscillator model to describe the cell-cycle phase and volume growth dynamics. Building on the experimental work of Charvin *et al.*, we devised an open-loop control strategy that relies on the concept of phase-locking and, supported by our preliminary experiments, demonstrated its efficacy in synchronising the cell-cycle across a population of cells. However, open-loop control strategy faces several hurdles in a practical setting and relies on the proper choice of the forcing period. To overcome such limitations, we devised a feedback control strategy based on Model Predictive Control, which has been already successfully applied to biological control problems. Through numerical analysis, we demonstrated the feasibility of using MPC to synchronise the cell-cycle phases, obtaining better performance when compared to the open-loop strategy. Nevertheless, several issues remain open both in terms of control performance and experimental implementation. Indeed, model uncertainty and parametric variability could affect the control performance, thus adaptive and robust control strategies may be required.

Moreover, estimation of the cell-cycle phase from the YFP fluorescence signal is non-trivial and requires improvement in image segmentation algorithms.

## Acknowledgements

This work was supported by the COSY-BIO (Control Engineering of Biological Systems for Reliable Synthetic Biology Applications) funded by the European Union’s Horizon 2020 research and innovation programme under grant agreement No 766840.

